# NoRCE: Non-coding RNA Sets Cis Enrichment Tool

**DOI:** 10.1101/663765

**Authors:** Gulden Olgun, Afshan Nabi, Oznur Tastan

**Affiliations:** Department of Computer Engineering, Bilkent University, Ankara, Turkey; Faculty of Engineering and Natural Sciences, Sabanci University, Tuzla, Istanbul, Turkey

## Abstract

**Summary:** While some non-coding RNAs (ncRNAs) are assigned to critical regulatory roles, most remain functionally uncharacterized. This presents a challenge whenever an interesting set of ncRNAs needs to be analyzed in a functional context. Transcripts located close-by on the genome are often regulated together. This genomic spatial proximity can lead to a functional association. Based on this idea, we present a tool, NoRCE, that performs *cis* enrichment analysis for a given set of ncRNAs. Enrichment is carried out using the functional annotations of the coding genes located proximal to the input ncRNAs. NoRCE allows incorporating other biologically relevant information such as topologically associating domain (TAD) boundaries, co-expression patterns, and miRNA target prediction information. NoRCE repository provides several data, such as cell-line specific TAD boundaries, functional gene sets, and expression data for coding and ncRNAs specific to cancer for the analysis. Additionally, users can utilize their custom data files in their investigation. Enrichment results can be retrieved in a tabular format or visualized in several different ways. NoRCE is currently available for the following species: human, mouse, rat, zebrafish, fruit fly, worm, and yeast. NoRCE is a platform-independent, user-friendly, comprehensive R package that could be used to gain insight into the functional importance of a list of any type of interesting ncRNAs. Users can run the pipeline in a single function; also, the tool offers flexibility to conduct the users’ preferred analysis in a single base and design their pipeline. It is available in Bioconductor and https://github.com/guldenolgun/NoRCE.

## 1 Introduction

The advent of next-gen sequencing technologies and their application to transcriptomes have shown that the vast majority of the human genome is transcribed [1, 2] and the non-coding RNAs(ncRNAs) represent the largest class of transcripts in the human genome [3, 4]. NcRNAs are categorized into different groups based on length, location, or function: long noncoding RNAs (lncRNAs), microRNAs (miRNAs), small interfering RNAs (siRNAs), small nucleolar RNAs (snoRNAs), small nuclear RNAs (snRNAs), and Piwi-interacting RNAs (piRNAs).

NcRNAs have been implicated in a wide array of cellular processes [5, 2, 6, 7] and emerging evidence further reinforces that they have crucial functional importance for normal development and disease [8]. For example, lncRNAs, the largest class of ncRNAs, are reported to control nuclear architecture and transcription, modulate mRNA stability, translation, and post-translational modifications [9, 7]. Nevertheless, only a small fraction of ncRNAs have been functionally characterized today, and functions of the majority of ncRNAs remain unknown. The lack of functional annotation of ncRNAs presents a challenge when an ncRNA set of interest is available and needs to be functionally investigated for further analysis.

Most of the available ncRNAs functional enrichment tools are limited to miRNAs. In the first step of these tools, they make a list of genes that are targeted by at least one of the miRNAs in the input set, which is followed by an enrichment analysis on this target gene set[10, 11, 12]. The target set is derived from experimentally validated interaction databases or produced by target prediction algorithms. Among them, Corna [10], miRTar [12], and Diana-miRPath v.3 [11] differ from varied features such as the source of the targets or the functional sets on which the analysis is conducted. Since the predicted target interactions might include high false positives and are not context-specific, some methods also take into account the changes in mRNA levels. MiRComb [13] conducts a miRNA-mRNA expression analysis followed by miRNA target prediction on the negatively correlated mRNA targets. miRFA [14] considers both the negatively and positively correlated using TCGA data. miTALOS [15, 16] additionally provides a tissue-specific filtering of the targets.

There is also a limited number of tools that offer functional annotation and enrichment analysis on lncRNA sets. Similar to miRNA methods, these methods first find a set of coding genes that are co-expressed genes with the given lncRNA or the lncRNAs in the collection and conduct analysis on these coding genes [17, 18, 19]. With regards to other ncRNAs, only a few studies provide analysis for ncRNAs other than lncRNA and miRNA. StarBase v2 first constructs a regulatory network based on experimentally identified RNA binding sites and their interactions; next, they perform functional enrichment on the interacting coding genes of the ncRNAs [20]. Starbase v2 offers analysis on miRNAs, lncRNAs, and the pseudogenes. CircFunBase [21] is not an enrichment tool but provides manually curated functions of circular RNAs that can be used for enrichment analysis.

The available tools are limited to the type of input they provide and do not take into account genomic neighborhood information. In this work, we present NoRCE (Non-coding RNA Sets Cis Enrichment Tool), which offers broad applicability and functionality for enrichment analysis of all types of ncRNAs sets using genomic proximity. NoRCE first finds nearby coding genes on the genome of the ncRNAs in the input set and uses the functional annotations of this coding gene set to perform functional enrichment on the ncRNA set. The motivation of this functional annotation transfer is based on the evidence presented earlier that genes nearby can be linked functionally. Thevenin et al. show that functionally related coding genes are co-localized on the genome [22]. Engreitz et al. [23] report that both coding and non-coding genes can regulate the expression of neighboring genes on the genome. There are several instances of lncRNAs that influence the nearby genes’ expressions [24, 25, 26]. For example, ÃŸrom et al. [27] report that the depletion of some ncRNAs led to decreased expression of their neighboring protein-coding genes. Others also support the involvement of lncRNAs in the *cis* regulation, where both the regulatory ncRNA and the target gene are transcribed from the same or nearby genomic locus [28]. Based on these findings, in this work, we take into account the coding genes nearby to functionally assess a given ncRNA set. The transfer of functional annotation from nearby coding genes has been used in the general genomic interval set enrichment tools [29, 30, 31, 32].

To offer broad functionality and applicability, NoRCE allows several additional features. The identified neighborhood coding gene set can be filtered or expanded with coding genes found to be co-expressed with the input ncRNAs. The co-expressed genes can be used to expand the set. For this, NoRCE allows users to input their expression data or make use of pre-computed correlation results for The Cancer Genome Atlas (TCGA) project expression data. Since TAD boundaries affect the expression of neighboring genes [33], NoRCE also allows analysis that takes into account the topologically associated regions (TAD) boundaries on the genome. NoRCE provides miRNA specific options as well; the user has an option to filter the neighbor set with predicted targets of the input miRNAs. Moreover, the input ncRNA set can be filtered based on ncRNA biotype (such as sense, antisense, lincRNA). NoRCE supports various commonly used statistical tests for enrichment.

In the following sections, we first detail the NoRCE’s capabilities and the technical details. We also exemplify the NoRCE on two different functional analyses. In the first use case, we analyze the set of ncRNAs differentially expressed in brain disorder, while the second one showcases miRNA specific analysis on cancer patient data.

## 2 Implementation

Capabilities of NoRCE and workflow are summarized in Figure 1. For a given set of ncRNAs, NoRCE first recognizes the coding genes close to ncRNA genes on the linear genome. Based on user-specified options, these genes are expanded or filtered using co-expressed genes, target predictions, or using the information on the TAD regions. Once the genes of interest are gathered, several gene enrichment analyses are performed. The details of these steps are provided in the following sections.

**Figure 1:**
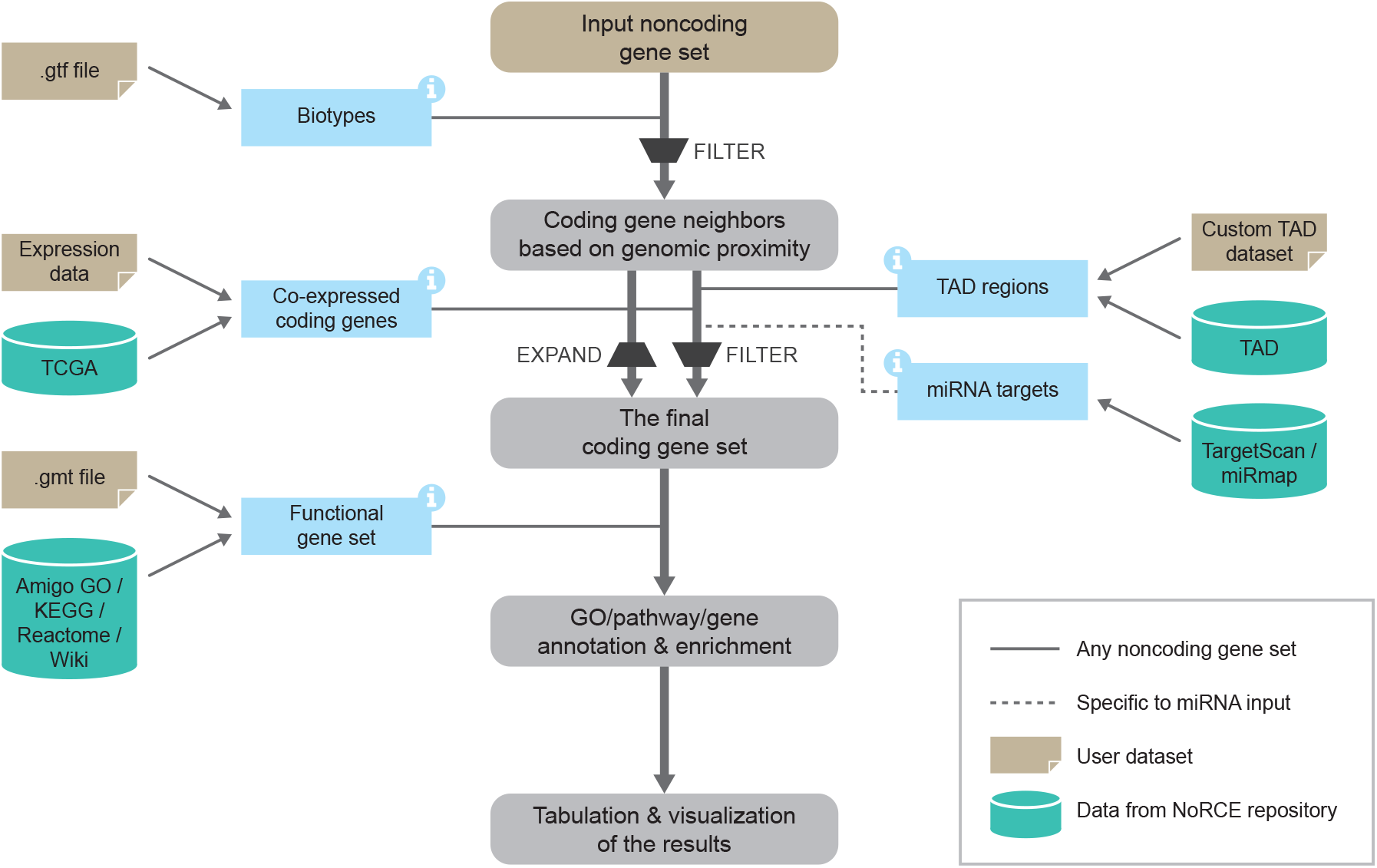
The workflow of the NoRCE package.

### 2.1 Species Supported

NoRCE supports analysis for *Homo sapiens, Mus musculus* (house mouse), *Rattus norvegicus* (brown rat), *Danio rerio* (zebrafish), *Drosophila melanogaster* (fruit fly), *Caenorhabditis elegans* (worm) and *Saccharomyces cerevisiae* (yeast). For *Homo sapiens*, it handles human hg19 and hg38 assemblies. For the other species, it uses the most recent assembly of the species. Supported assemblies for different species are provided in Supplementary Files1 Table S1.

### 2.2 Curating the *cis* Coding Gene List

NoRCE accepts a set of any type of ncRNAs, *S* = {*r*_1_, …, *r*_*n*_}. For each ncRNA, *r*_*i*_ ∈ *S*, in the input list, NoRCE identifies all spatially proximal protein-coding genes. The proximal genes are considered as those that are within the base-pair limit of the genomic start coordinate of the input gene and/or within the base-pair limit of the genomic end coordinate of the input gene. If the coding gene *r*_*i*_ is located within the user-specified base-pair limit from the upstream and/or downstream of known transcription start and/or end position of the ncRNA gene, it is designated as a neighboring coding gene of *r*_*i*_ and added to the coding gene list pool of *C*_*i*_. The union of the coding genes, constitute the final coding gene set to be tested for functional enrichment, 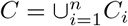. The pool of coding genes can be further filtered or expanded based on the additional biological evidence available, detailed in the next sections, and user-selected options. Users can also limit the analysis to the introns or exons of the neighboring coding genes. In that case, NoRCE applies the genomic proximity criterion on the intron or the exon of the genes based on the user’s selection.

Gene coordinates and their annotations are retrieved from two different databases: ENSEMBL [34] and UCSC [35] since no single source contains information on all the transcripts. We collect the ENSEMBL data via biomaRt package [36]. Genes are retrieved from UCSC using the rtracklayer package[37].

### 2.3 Co-expressed Non-coding Genes

Since coding genes that exhibit high co-expression patterns can hint to functional cooperation, NoRCE enables the user to incorporate co-expressed coding genes into the analysis. If the filtering option is set, each *C*_*i*_ is filtered such that only the neighboring coding genes that are also co-expressed with *r*_*i*_ are placed into *C*. If the expansion option is set, a coding gene is co-expressed with any of the *r*_*i*_∈ *S* is added to *C*.

NoRCE enables the user to conduct the expression analysis with user input expression data. In this case, users are expected to load the expression data in TSV or TXT format; or they can use the *SummarizedExperiment* object in R. Before the correlation analysis, NoRCE executes a pre-processing step on expression data. The variance of each gene’s expression is calculated, and genes that vary lesser than the user-defined variance cutoff, 0.0025 by default, are excluded from the analysis. NoRCE supports commonly used correlation measures: Pearson Correlation, Kendall Correlation, and Spearman Correlation. The default values for correlation coefficient cutoff is 0.3, for significance p-value, 0.05 and confidence level 0.95. The user can set the correlation and significance cutoffs based on their need.

To assist analysis for cancer, NoRCE also allows using pre-computed co-expressed gene sets for ncRNAs measured in The Cancer Genome Atlas (TCGA) project [38]. Since TCGA contains the expression profiles for miRNA, mRNA, and lncRNA, this examination is limited to only the miRNA and lncRNA inputs. The co-expressed genes are defined using the Pearson correlation coefficient. Users can set the cutoff for the correlation coefficient.

### 2.4 Filtering Genes With the TAD Boundary Information

The gene regulatory interactions are affected by the 3D chromatin structure of the genome [39]. On a single chromosome, chromatin compartmentalizes into sub-domains, named as topologically associating (TADs). TAD boundariesâĂŤregions insulate the *cis*-regulating elements [40]. NoRCE allows filtering based on TADs. If this option is selected, when curating the nearby genes of an ncRNA, NoRCE will only include the coding genes within the same TAD boundary with that of the ncRNA. We compile TAD regions for different cell-lines and species from various sources and made them available for use in conducting the analysis. These data sources and the species they are available for is provided in Supplementary File1 Table S2. NoRCE allows inputting BED formatted TAD boundary files. Thus, the user can conduct this analysis with other available TAD information.

### 2.5 Biotype Specific Analysis

If the user wants to conduct a biotype specific analysis, NoRCE can select the ncRNAs that are annotated with the given biotypes and use this biotype-filtered subset in the subsequent steps. Also, NoRCE allows us to extract the given biotypes from *S* and perform analysis on the subset of genes that do not contain the genes annotated with given biotypes. NoRCE accepts GTF formatted GENCODE annotation files for biotype analysis.

### 2.6 miRNA Target List

For miRNA specific inputs, NoRCE provides additional features. The coding gene set, *C*, can be restricted to the potential miRNA targets; thus, only neighboring coding genes that are also miRNA potential targets are included. The miRNA target list is curated from various sources. Computationally predicted miRNA-target interactions are obtained from the TargetScan [41] for the species except *Rattus norvegicus* as it is not available. Target predictions for *Rattus norvegicus* miRNAs are obtained from the miRmap [43]. No miRNA is reported for *Saccharomyces cerevisiae* [46]. Thus, NoRCE does not provide any miRNA analysis for *Saccharomyces cerevisiae*. Table 1 presents the details of the pre-computed target predictions.

**Table 1:** List of miRNA target prediction algorithms used for each species. The version information is provided, which is the current available version for the corresponding species.

| Species | Database | Ver/Date | Ref. |
| --- | --- | --- | --- |
| *Homo sapiens* | TargetScan | v7.2 | [41] |
| *Mus musculus* | TargetScan | v7.1 | [42] |
| *Rattus norvegicus* | miRmap | 2013 | [43] |
| *D. melanogaster* | TargetScan | v6.2 | [44] |
| *Danio rerio* | TargetScan | v6.2 | [42] |
| *C. elegans* | TargetScan | v6.2 | [45] |

### 2.7 Enrichment Analysis

Once the coding gene list, *C*, is curated, NoRCE conducts functional enrichment analysis. NoRCE supports analysis with various functional annotations: gene ontology (GO), Kyoto Encyclopedia of Genes and Genomes (KEGG), Reactome pathway, WikiPathways, genes, or GMT formatted integrated pathway dataset. For the annotation, we make use of biannually updated databases in Bioconductor. Gene ontologies and their annotated gene list are provided via GO.db [47] package. To increase the statistical analysis power, only the GO terms with at least 5 annotated protein-coding genes are considered as suggested by [17]. KEGG annotation is performed using KEGG.db [48], for Reactome enrichment analysis, NoRCE utilizes reactome.db [49]. NoRCE employs WikiPathways API to retrieve the pathways and annotated gene list [50].

NoRCE supports commonly used enrichment tests: hypergeometric test, Fisher Exact Test, Binomial Test, *X*^2^ test. We refer readers to [51] for details of the statistical tests. The background gene set is all the genes in the functional annotation source that is selected. NoRCE provides the flexibility of providing a user-defined background gene set.

### 2.8 Presentations of the Results

NoRCE provides different ways to export the results. All information in enrichment analysis can be retrieved in a tabular format. Also, users can set the number of top enrichment results to exported, and NoRCE outputs these results based on p-value or p-adjusted values in a tabular format. Networks and dot plots can be used to visualize the enrichment results. The dot plot shows the top enriched terms, their p-values (or p-adjusted values), and the number of enriched genes in the input neighbor set. In the network representation, the enriched terms are represented with nodes, and the ncRNA and coding transcripts related to the enriched terms are represented with edges. In this graph, the node size is proportional to the node degree. The nodes in the networks are clustered, and a color code distinguishes between node clusters. Modularity clustering is employed to cluster the nodes [52]. For network visualization features, NoRCE makes use of the igraph package [53].

NoRCE also offers specialized visualization options for pathway and GO analysis. GO enrichment results can be illustrated in a directed acyclic graph (GO-DAG). We derive the DAG information through the AmiGO API [54]. In this diagram, nodes are GO-terms, and edges indicate relation types between GO terms. Enriched GO-terms are colored according to their p-values or p-adjusted values. Users can export enriched GO-DAG diagrams in a PNG or SVG format. For pathway enrichment results, KEGG and Reactome enrichment results can be visualized within KEGG and Reactome maps. The enriched terms are marked with color using the KEGG and Reactome APIs, respectively. These visuals are displayed through the browser. In the results sections and the supplementary materials, we provide examples of these visualizations.

## 3 Results

To demonstrate how NoRCE could be used to analyze a list of ncRNAs functionally, we apply NoRCE on several problems and datasets. We use the default parameter settings in the following analyses unless otherwise stated.

### 3.1 Case Study 1: Enrichment Analysis of the ncRNAs for the Psychiatric Disorders

In this use case, we demonstrate the functional enrichment analysis of a set of ncRNAs related to brain disorders based on gene expression data measured by Gandal et al. [55]. These ncRNAs exhibit gene- or isoform-level differential expression in at least one of the following disorders: autism spectrum disorder (ASD), schizophrenia (SCZ), and bipolar disorder (BD). In total, the ncRNA gene set contains 1,363 differentially expressed human ncRNAs. We perform GO enrichment for biological processes and pathway enrichment analysis based on pathways provided by Bader Lab [56]. The number of pathways and the different pathway sources included in the Bader Lab set is provided in Supplementary File 1 Table S3. In these enrichment analyses, the background gene sets are described as the groups of all annotated genes in the corresponding GO or pathway dataset. The protein-coding genes that fall into this neighborhood region of the ncRNAs are input to the enrichment analysis. We also showcase NoRCEâĂŹs ability to constrain the input set with protein-coding genes within the TAD boundaries.

#### 3.1.1 Functional enrichment results

The dot plot in Figure 2 shows the top 35 enriched BP GO-terms, sorted based on the significance of enrichment. The number of annotated genes with the corresponding GO-terms are provided in the graph. We detect RNA related GO terms such as the *positive regulation of pri-miRNA transcription by RNA polymerase II, miRNA mediated inhibition of translation*, and *RNA processing*. Additionally, there GO terms which are pertinent to various neurological functions such as *response to ischemia, sensory perception of pain*, and *neurogenesis*. It has been reported that cerebral ischemia-induced genes are upregulated in schizophrenia [57], and it is common to have chronic pain in bipolar patients [58]. Interestingly, we observe that the enriched terms include cardiac and vascular-related functions. Several studies exhibit interactions between neural diseases and changes in blood vessel pathology and blood flow [59, 60, 61]. Others reveal that patients with bipolar disorder have low heart rate variability, which is a physiological measure of variation in the time interval between each heartbeat [62, 63].

**Figure 2:**
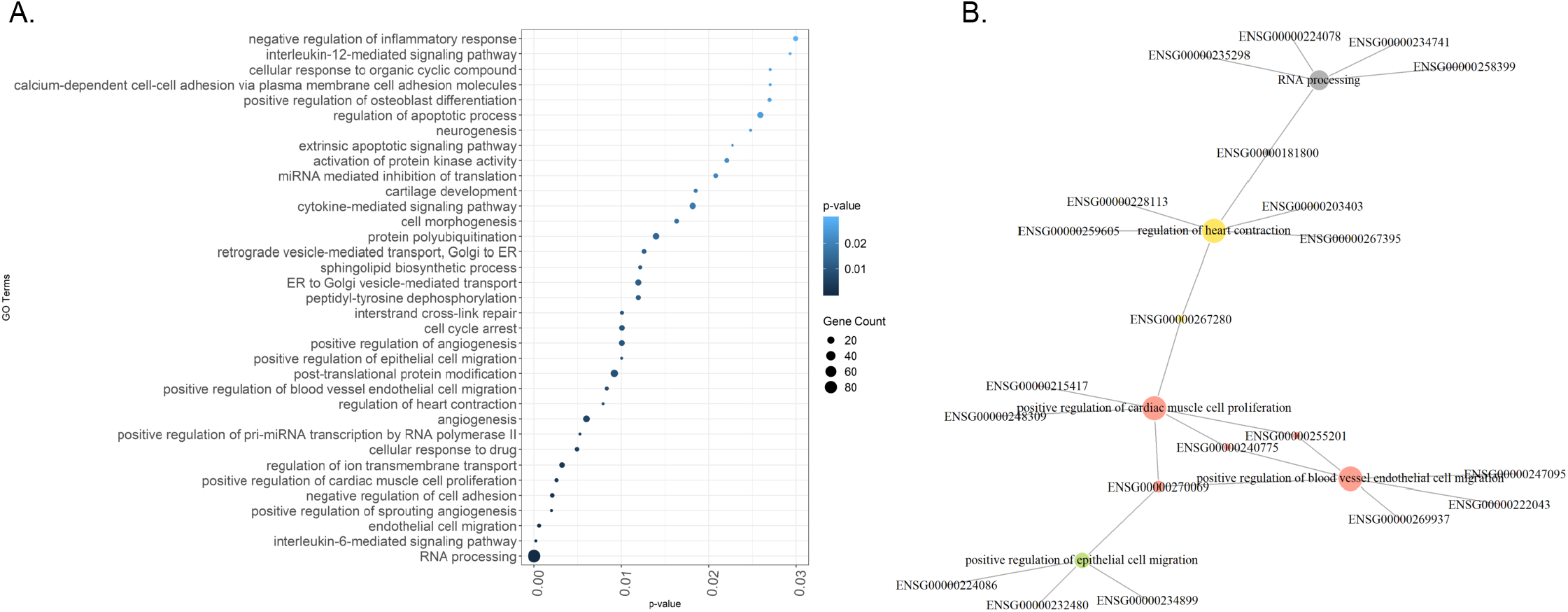
A) Top 35 GO biological processes enriched in the set of differentially expressed brain ncRNAs from [55]. The x-axis represents the p-values; the y-axis represents the GO terms. The dot area is proportional to the size of the overlapping gene set, and the color signifies the p-value of the enrichment test for the corresponding GO-term. B) The functional enrichment repeated with TAD filtering GO-term:ncRNA network. The size of the nodes is proportional to the degree of the node. Different colors are used to differentiate the clusters of nodes.

Alternative visualizations of these functional enrichment results are provided in the Supplementary Materials section.

Supplementary Notes Figure S1 shows the top 7 GO terms in a network visualization format. Supplementary Notes Table S4 lists the top enriched GO BP terms to showcase the tabular format output capabilities.

#### 3.1.2 Functional Enrichment Results with TAD Filtering

We repeat the previous functional analysis when the TAD filtering is on. When this filter is applied, only the protein-coding genes near the ncRNAs in the input list, and at the same time reside within the same TAD regions are included in the enrichment analysis. In this analysis, we use custom defined TAD regions for the adult dorsolateral pre-frontal cortex that are provided by the [55] study, and we keep all the other parameters in their default values.

Fig 2 illustrates the GO-term network for top 7 enriched GO terms. Alternative representations of these results are provided in the Supplementary Materials (Supplementary Notes Table S5, Figures S2 (A), and S2 (B)). We recognize similar functional patterns with the previous analysis. We notice brain-related GO-terms, including central nervous system development and *calcium ion transport*. Also, several cardiac and vascular-related functions are observed (*positive regulation of cardiac muscle cell proliferation, positive regulation of blood vessel endothelial cell migration* and *angiogenesis*). Different from the previous analysis, we identify cell cycle regulation related to GO-terms. Cell cycle regulating genes have been associated with autism in GWAS studies, and in DNA derived from the pre-frontal cortex, they showed autism-specific CNVs [64]. In schizophrenia and bipolar disorder, many genes involved in cell cycle regulation have been shown to have differential expression levels [65]. Cell adhesion has been reported to be disrupted in autism [66]. In schizophrenia, cell adhesion regulating genes are dysregulated, too [67]. Moreover, in schizophrenia and bipolar disorders, cell adhesion pathways have been reported to contribute to disease susceptibility [68].

#### 3.1.3 Pathway enrichment using predefined pathway gene sets

NoRCE enables pathway enrichment analysis for various sources, including KEGG, Reactome pathway, and WikiPathways. Also, NoRCE supports pathway enrichment using custom pathway databases such as MSigDb [56], or other user-curated data provided in GMT format. To showcase the NoRCE capability, we utilized the Bader Lab dataset as the user-defined pathway gene set analysis. In this analysis, we only consider the genes in the neighborhood of the differentially expressed ncRNAs in brain disorders.

Interestingly, we find many enriched pathways that are related to neural diseases. Some of these pathways directly related to ASD, schizophrenia, and bipolar disorder, including `Synaptic signaling pathway` associated with an `autism spectrum disorder, WP4539, Amyotrophic lateral sclerosis, WP2447, Alzheimer’s disease, WP2059`. Additionally, we found that many of the signaling pathways, such as *G-Protein Signaling, mTOR signalling, MAPK Signaling* pathway and those pathways are associated with at least one of the brain disorder: autism spectrum disorder, schizophrenia, and bipolar disorder [69, 70, 71]. Due to the space limit, we list a subset of enriched disease-related pathways in Supplementary Table S6 and S7.

#### 3.1.4 Comparison Between ASD Associated GO-terms and NoRCE Enrichment Results

Krishnan et al. [72] predict novel ASD risk genes based on the brain-specific functional network. In their work, they also identified functions potentially dysregulated by ASD-associated mutations. We compare our findings with this set of ASD associated GO function terms [72]. We observe that most of the enriched GO terms reported in NoRCE are listed as potential ASD-related GO terms in Krishnan et al. [72] study. The ncRNA enrichment analysis of NoRCE without TAD filtering identifies 48 enriched GO terms, and 32 of these terms are also in the list of ASD related GO terms [72], corresponding to 67% overlap. When TAD filtering is applied, there were 29 enriched terms, and 22 were in the ASD GO term list, corresponding to 75% overlap (Supplementary Table S9). We tested the significance of these overlap ratios. We randomly selected ncRNA set with the same size of the input from the all gene population. We found the enriched term with this random gene set and checked if the overlap ratio is equal or higher in the randomized case. A *p*-value is calculated by repeating this procedure 1, 000 times.

### 3.2 Case Study 2: Functional enrichment analysis of variably expressed miRNAs in brain cancer using miRNA targets

NoRCE offers a filtering option for the input miRNA’s targets. Users can choose to filter ncRNA neighbors, such that only those that are the targets of the miRNA are included. To demonstrate this option, we use a set of miRNAs that are differentially expressed ncRNAs in brain cancer obtained from dbDEMC 2.0 [73] for the functional analysis. This set contains 407 miRNAs and is provided in NoRCE with the name *brain_miRNA*. We choose the Reactome pathway as the functional gene set.

We identify *lysosome vesicle biogenesis* (*p*-value = 7.1e-05), *trans-golgi network vesicle budding* (*p*-value = 0.0006), *ion channel transport* (*p*-value= 0.0091), and *axon guidance* (*p*-value = 0.0382) pathways as enriched. Previous studies report that the *axon guidance* and *ion channel transport* pathways are related to the Glioblastoma Multiforme [74, 75]. Other evidence also suggests that miRNAs could be acting as key fine-tuning regulatory elements in *axon guidance* [76].

### 3.3 Case Study 3: Functional enrichment analysis with co-expression analysis

NoRCE also supports filtering based on a co-expression analysis. When defining coding gene neighborhoods for an ncRNA, the user can choose to include a coding gene only if it is coexpressed. Alternatively, the users can choose to augment the coding genes list with the co-expressed coding gene set. To demonstrate this option, we use NoRCE on the brain cancer patient data obtained from TCGA.

The TCGA data include expression levels for mRNA and miRNA for matched primary tumor solid samples from 527 tumor patients. miRNA-seq data are measured as per million mapped reads (RPM) values, and RNA-seq data are measured as Fragments per Kilobase of transcript per Million mapped reads upper quartile normalization (FPKM–UQ). We apply the same pre-processing step as in our previous method [77]. Genes and miRNAs that have very low expression levels (RPKM *<* 0.05) in many patients (more than 20% of the samples) are filtered out. The gene expression values are *log*2 transformed, and those with high variability are retained for co-expression analysis. For this aim, only the genes with median absolute deviation (MAD) above 0.5 are used. The final expression dataset contains 444 miRNA and 12, 643 mRNA genes on 527 tumor patients on which we perform Pearson correlation analysis. The mRNAs which have more than 0.1 correlation with a miRNA are retained.

When we examine the enriched pathways, *Signaling by Receptor Tyrosine Kinases* emerges as an important pathway. *Receptor Tyrosine Kinases* is a cell surface receptor family, and its members are responsible for growth factors, hormones, cytokines, neurotrophic factors [78]. Following their activation, they can signal through downstream pathways responsible for survival, differentiation, and angiogenesis [78]. Inhibition on *Receptor Tyrosine Kinases* and their signal pathways are utilized as target therapy on brain cancer [79]. Also, miRNAs are reported to take a role as mediators or suppressors in these pathways and promote tumor cell death [79]. The Reactome diagram for this pathway is illustrated in Supplementary Notes Figure S4. In both target-based and co-expression-based analyses on the differentially expressed miRNAs, *Axon guidance pathway* is enriched. MiRNAsâĂŹ role in axon guidance have been reported elsewhere [76, 80, 81, 82]. For example, Baudet et al. [82] report that miR-124 controls Sema3A, which is essential for normal axon guidance. Accumulating evidence also points out that axonogenesis is stimulated by malignant cells and contributes to cancer growth and metastasis [83]. These findings also support the NoRCE capability for finding interesting functional inferences. The details of the enrichment results are provided in Table 2.

**Table 2:** Pathway enrichment results for nearby co-expressed genes with miRNAs. The GeneRatio is computed by dividing the overlapping with the coding genes with the functional gene set to the number of all protein-coding genes within the input set neighbourhood. The BGRatio column represents the ratio of the number of genes found in the enriched GO term set to the size of the background gene set. The EGNo refers to the size of the overlap between the corresponding GO term gene set and the neighboring coding gene set. ncGeneList column contains ncRNA genes that are enriched with the corresponding GO-term.

| Pathway ID | Pathway Term | Pvalue | GeneRatio | BGRatio | EGNo | ncGeneList |
| --- | --- | --- | --- | --- | --- | --- |
| R-HSA-9006934 | Signaling by receptor tyrosine kinases | 0.0006 | 8/44 | 473/10654 | 8 | hsa-mir-199b<br>hsa-mir-214<br>hsa-mir-3934<br>hsa-mir-455<br>hsa-mir-483<br>hsa-mir-141<br>hsa-mir-338<br>hsa-mir-3689a |
| R-HSA-422475 | Axon guidance | 0.0017 | 8/44 | 553/10654 | 8 | hsa-mir-199b<br>hsa-mir-214<br>hsa-mir-4684<br>hsa-mir-585<br>hsa-mir-95<br>hsa-mir-150<br>hsa-mir-3689a |
| R-HSA-1266738 | Developmental biology | 0.0327 | 9/44 | 1097/10654 | 9 | hsa-mir-199b<br>hsa-mir-214<br>hsa-mir-4684<br>hsa-mir-585<br>hsa-mir-935<br>hsa-mir-95<br>hsa-mir-150<br>hsa-mir-3689a |

## 4 Discussion and Conclusion

NoRCE is a comprehensive, flexible, and user-friendly tool for enrichment analysis of all types of ncRNAs. It works for multi-species and is available as an R package. NoRCE, unlike existing tools, conducts enrichment by taking into account the genomic neighborhood of the ncRNAs in the input set and transfer functional annotations of these coding genes. We should note that though although cis-regulation has been reported for many ncRNA types, it may not hold for all types of ncRNAs. Therefore, in addition to the genomic neighborhood-based analysis, NoRCE allows the standard approaches of using coding genes co-expressed with the input ncRNAs in detecting the enriched functions. Another unique feature of NoRCE that it allows an option for making use of TAD regions. NoRCE provides flexibility to the user; the user can analyze with different options and use the library’s readily available datasets to conduct the analysis or input custom datasets. Also, she can include or exclude any analysis that NoRCE contains.

In showcasing NoRCE, we analyzed ncRNAs obtained in different studies, including brain disorder related ncRNAs and cancer-related datasets. These analyses yield interesting biological findings highlighting how NoRCE could be useful in answering a wide range of questions. The datasets and examples are also provided in the R*\*Bioconductor package.

NoRCE uses functional sets such derived from GO and pathway databases and miRNA prediction tools. Improvements in these databases and tools will help NoRCE to make a more accurate analysis. NoRCE is designed for non-coding RNAs, but one can use both coding and non-coding RNAs as input. Currently, the user can use NoRCE to conduct analysis in human and mouse, rat, zebrafish, fruit fly, worm, and yeast. As a future direction, NoRCE can be extended to support analysis for other species. Moreover, the current version of the package contains only miRNA target predictions. However, we can enhance our target prediction parts for other functional ncRNAs, including sRNAs and snoRNAs.

## Author’s contributions

G.O and O.T. designed the study. All authors contributed to the results, discussions, and writing of the manuscript.

## Acknowledgements

We thank Sabanci University and Bilkent University for internal funding support. The results shown here are in part based upon data generated by the TCGA Research Network: https://www.cancer.gov/tcga. We also thank Marcel Ramos, our R Bioconductor package code reviewer, Martin Morgan, Michael Lawrence, and Lori Shepherd for their valuable feedback, comments, and time on our R \Bioconductor package.

