## Supplementary File1 for "NoRCE: Non-coding RNA Sets Cis Enrichment Tool"

### 1 Supplementary Information on Methods

#### 1.1 Data Sources

NoRCE repository contains various datasets. Below, we detail these sources.

##### 1.1.1 Gene and Gene Ontology Annotations

Table S1: The supported assemblies for different species in NoRCE.

| Species | Supported Assembly | |
|---|---|---|
| | UCSC | NCBI |
| *Homo sapiens* | hg19 | GRCh37, GCA_000001405.1, Feb. 2009 |
| *Homo sapiens* | hg38 | GRCh38, GCA_000001405.15, Dec. 2013 |
| *Mus musculus* | mm10 | GRCm38.p6, INSDC Assembly GCA_000001635.8, Jan 2012 |
| *Rattus norvegicus* (brown rat) | rn6 | Rnor_6.0, INSDC Assembly GCA_000001895.4, Jul 2014 |
| *Drosophila melanogaster* (fruit fly) | dm6 | BDGP6, INSDC Assembly GCA_000001215.4, Jul 2014 |
| *Danio rerio* (zebrafish) | danRer10 | GRCz10, GCA_000002035.3, Sep. 2014 |
| *Caenorhabditis elegans* (worm) | ce11 | WBcel235, GCA_000002985.3, Feb. 2013 |
| *Saccharomyces cerevisiae* (yeast) | sacCer3 | UCSC version sacCer3, 2011 |

##### 1.1.2 TAD Boundaries Maintained by NoRCE

Table S2: Topological associating domain data that included in the NoRCE repository

| Species | Name | Ver. | # of cell lines | # of TAD regions | Source |
|---|---|---|---|---|---|
| *Homo sapiens* | tad_hg19 | hg19 | 37 | 74,424 | 3D Genome Browser[5] |
| *Homo sapiens* | tad_hg38 | hg38 | 42 | 96,526 | 3D Genome Browser[5] |
| *Mus musculus* | tad_mm10 | mm10 | 5 | 16,866 | 3D Genome Browser [5] |
| *D. melanogaster* | tad_dmel | dm6 | 1 | 2,846 | HiCBrowser[3] |

##### 1.1.3 Details of the Custom Pathways

#### 1.2 Data in NoRCE repository for the vignette and supplementary results

- brain_disorder_ncRNA : A list of ncRNAs differentially expressed in three psychiatric disorders including autism spectrum disorder (ASD), schizophrenia (SCZ), and bipolar disorder (BD) generated by Gandal et al. [2]. The set comprises1,363 ncRNAs. (Case study 1, Section 3.1).
- tad_custom : The TAD regions for dorsolateral prefrontal cortex obtained from [2]. The dataset contains 2,735 TAD regions. (Case study 1, Section 3.1).

Table S3: The number of pathways and the different pathway sources included in the January 2020 Bader Lab pathway data set.

| Pathway Database | Number of Pathways |
|---|---|
| NetPath | 25 |
| IOB | 33 |
| Panther | 173 |
| NCI | 223 |
| Cyc | 240 |
| MSigdb | 520 |
| WikiPathways | 559 |
| Reactome | 2,302 |
| Total | 4075 |

- miRCancerdb: The pre-computed Pearson correlation values of the expressions of pairs of $18,069,409$ miRNAs and genes for 34 cancer types. The data incorporates $1,015$ miRNAs, which are curated based on patient expression profiles that are provided by the TCGA. miRCancerdb is available at `https://figshare.com/articles/miRCancer_db_gz/5576329` [1]. (Case study 2, Section 3.2).

- brain_mirna : 407 differentially expressed miRNAs in human brain obtained from dbDEMC 2.0. (Case study 2, Section 3.2).

- breastmRNA : 667 differentially expressed mRNAs in human breast cancer. The analyze is conducted on patient expression data obtained from the TCGA project, collected on July $15^{th}$ 2017. Differential expression analysis is applied for $p-value < 0.05$ and $FDR < 0.05$ using Bioconductor limma package[4].

- mirna & mrna : Subset of brain cancer expression levels for mRNA, and miRNA obtained from TCGA. Data contains 527 matched tumor patients for 150 mRNA and 183 miRNA. Those datasets are subset of the pre-processed miRNA and mRNA expression data that utilized in Case Study 2 and they are intented to use for examples.

- ncRegion : The relevant regions of the differentially expressed human ncRNA genes (except pseudogenes) for three psychiatric disorders. The diseases include autism spectrum disorder (ASD), schizophrenia (SCZ), and bipolar disorder (BD), and the data were generated by Gandal et al. [2]. The final set contains 930 gene regions.

### 2 Supplementary Information on Results

#### 2.1 Case Study 1: Functional enrichment analysis of ncRNAs differentially expressed in psychiatric disorders

##### 2.1.1 Functional enrichment results

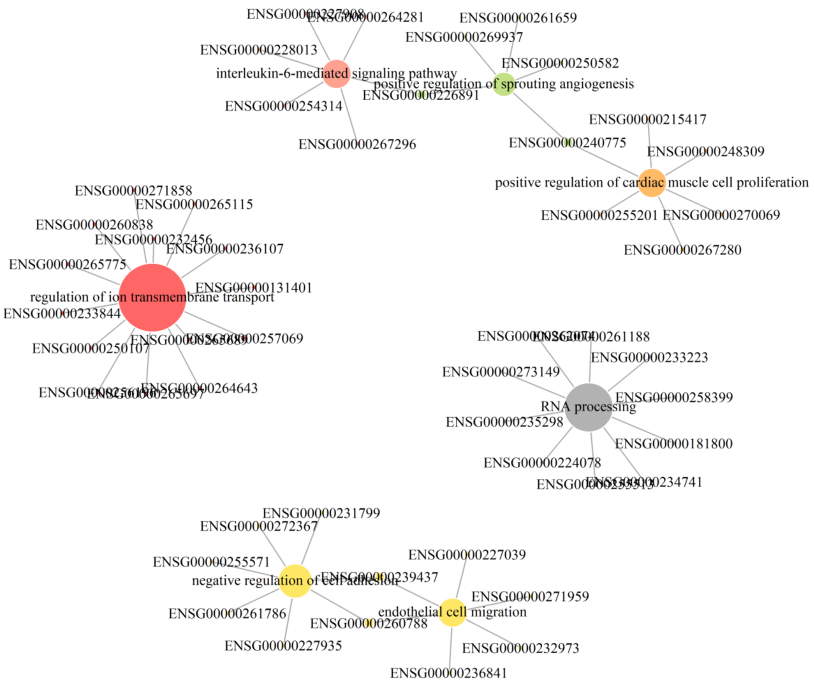

Figure S1: The top 7 enriched GO terms and the ncRNA genes which are associated with them are depicted as a network. The size of the nodes represents the degree, number of edges that are incident to the vertex, and each color represents different clusters in the network.

Table S4: Top 10 enrichment results of brain related biological process GO-terms that are obtained from the neighbourhood coding genes of ncRNA set of the [2]. The GeneRatio is computed by dividing the overlapping with the coding genes with the functional gene set to the number of all protein-coding genes within the input set neighbourhood. The BGRatiao column represents the ratio of the number of genes found in the enriched GO term set to the size of the background gene set. The EGNo refers to the size of the overlap between the corresponding GO term gene set and the neighboring coding gene set. ncGeneList column contains ncRNA genes that are enriched with the corresponding GO-term.

| ID | Term | Pvalue | GeneRatio | BGRatio | EGNo | ncGeneList |
|---|---|---|---|---|---|---|
| GO:0006396 | RNA processing | 1.984e-30 | 86/692 | 552/18671 | 86 | ENSG00000181800 ENSG00000235298 ENSG00000258399 ENSG00000224078 ENSG00000234741 ENSG00000262074 ENSG00000233223 ENSG00000255513 ENSG00000261188 ENSG00000273149 |
| GO:0070102 | interleukin-6-mediated signaling pathway | 2.142e-4 | 5/692 | 16/18671 | 5 | ENSG00000228013 ENSG00000254314 ENSG00000264281 ENSG00000226891 ENSG00000227908 ENSG00000267296 |
| GO:0043542 | endothelial cell migration | 5.806e-4 | 6/692 | 29/18671 | 6 | ENSG00000227039 ENSG00000232973 ENSG00000236841 ENSG00000239437 ENSG00000260788 ENSG00000271959 |
| GO:1903672 | positive regulation of sprouting angiogenesis | 1.976e-3 | 5/692 | 25/18671 | 5 | ENSG00000240775 ENSG00000250582 ENSG00000269937 ENSG00000226891 ENSG00000261659 |
| GO:0007162 | negative regulation of cell adhesion | 2.065e-3 | 7/692 | 49/18671 | 7 | ENSG00000227935 ENSG00000231799 ENSG00000239437 ENSG00000255571 ENSG00000260788 ENSG00000272367 ENSG00000261786 |
| GO:0060045 | positive regulation of cardiac muscle cell proliferation | 2.544e-3 | 6/692 | 38/18671 | 6 | ENSG00000215417 ENSG00000240775 ENSG00000248309 ENSG00000255201 ENSG00000267280 ENSG00000270069 |
| GO:0034765 | regulation of ion transmembrane transport | 3.165e-3 | 11/692 | 113/18671 | 11 | ENSG00000233844 ENSG00000236107 ENSG00000250107 ENSG00000256196 ENSG00000257069 ENSG00000264643 ENSG00000265115 ENSG00000265689 ENSG00000265697 ENSG00000265775 ENSG00000271858 ENSG00000131401 ENSG00000232456 ENSG00000260838 |
| GO:0035690 | cellular response to drug | 4.928e-3 | 7/692 | 57/18671 | 7 | ENSG00000224086 ENSG00000225555 ENSG00000248309 ENSG00000262251 ENSG00000269694 ENSG00000260549 ENSG00000260912 |
| GO:1902895 | positive regulation of pri-miRNA transcription by RNA polymerase II | 5.262e-3 | 5/692 | 31/18671 | 5 | ENSG00000184068 ENSG00000232480 ENSG00000250582 ENSG00000262251 ENSG00000266352 ENSG00000266936 |
| GO:0001525 | angiogenesis | 5.989e-3 | 17/692 | 232/18671 | 17 | ENSG00000226252 ENSG00000227694 ENSG00000229153 ENSG00000232480 ENSG00000232973 ENSG00000236841 ENSG00000237512 ENSG00000239437 ENSG00000254481 ENSG00000255201 ENSG00000258168 ENSG00000268854 ENSG00000269124 ENSG00000269386 ENSG00000227398 ENSG00000267280 ENSG00000273129 ENSG00000273374 |

##### 2.1.2 Filtering the close-by genes according to TAD boundaries

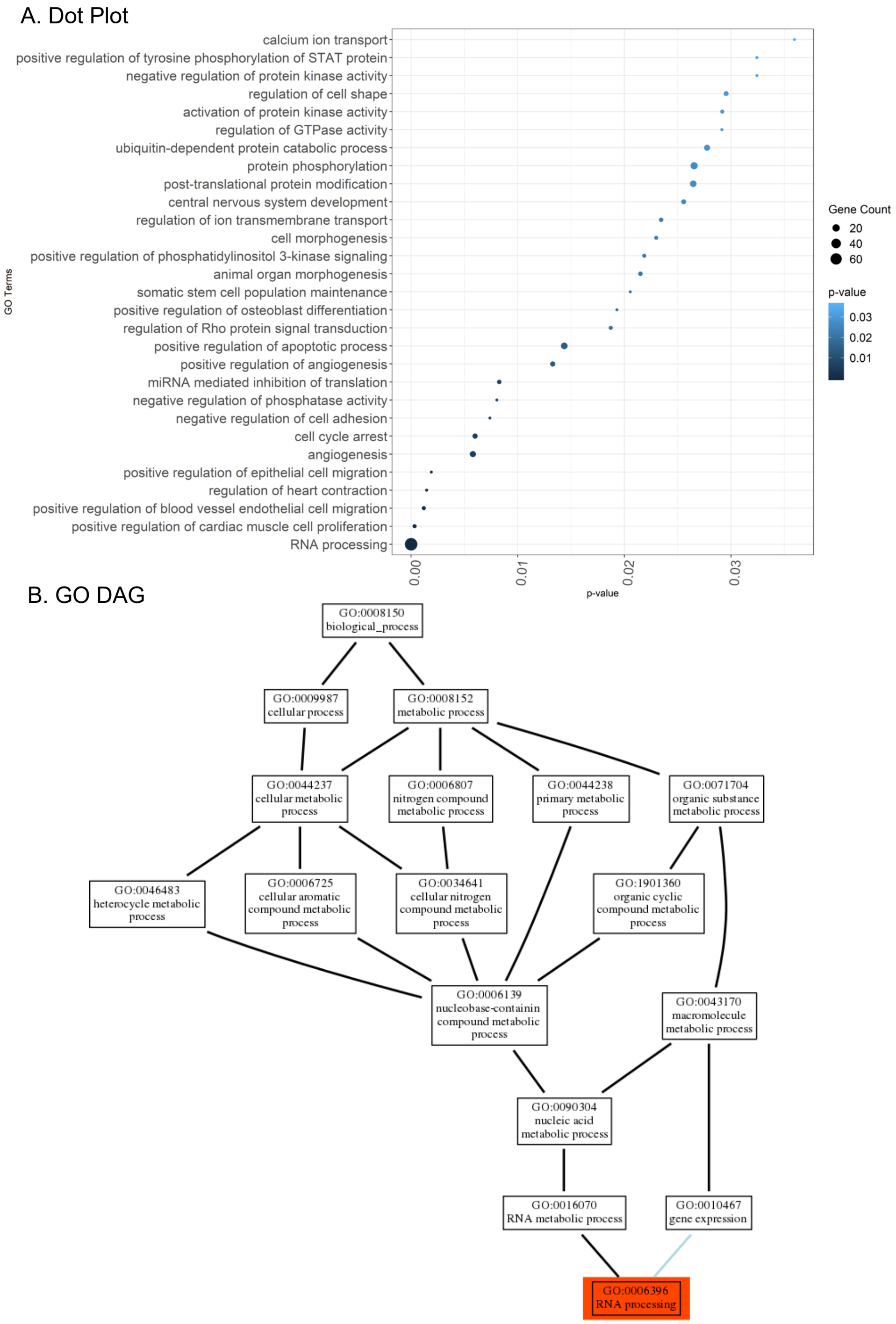

Figure S2: GO enrichment results for the genes that are in the same TAD boundaries with their close-by non-coding RNAs are only used. A) Top 35 GO biological process enrichment results pertaining to differentially expressed brain ncRNAs. The x-axis represent the *p*-value; the y-axis shows the GO term. The dot area is proportional to the size of the overlapping gene set, and the color signifies the *p*-value of the enrichment test for the corresponding GO-term. B) GO DAG diagram for the most enriched GO term. Enriched GO- terms are colored according to the p-value.

Table S5: Top 10 biological process GO term enrichment results when TAD filtering is used. The TAD information is from adult dorsolateral prefrontal cortex data [2] study. The GeneRatio is computed by dividing the overlapping with the coding genes with the functional gene set to the number of all protein-coding genes within the input set neighbourhood. The BGRatio column represents the ratio of the number of genes found in the enriched GO term set to the size of the background gene set. The EGNo refers to the size of the overlap between the corresponding GO term gene set and the neighboring coding gene set. ncGeneList column contains ncRNA genes that are enriched with the corresponding GO-term.

| ID | Term | Pvalue | GeneRatio | BGRatio | EGNo | ncGeneList |
|---|---|---|---|---|---|---|
| GO:0006396 | RNA processing | 1.417e-36 | 78/467 | 552/18671 | 78 | ENSG00000181800 ENSG00000235298 ENSG00000258399 ENSG00000224078 ENSG00000234741 |
| GO:0060045 | positive regulation of cardiac muscle cell proliferation | 3.179e-4 | 6/467 | 38/18671 | 6 | ENSG00000215417 ENSG00000240775 ENSG00000248309 ENSG00000255201 ENSG00000267280 ENSG00000270069 |
| GO:0043536 | positive regulation of blood vessel endothelial cell migration | 1.195e-3 | 6/467 | 48/18671 | 6 | ENSG00000222043 ENSG00000240775 ENSG00000247095 ENSG00000255201 ENSG00000269937 ENSG00000270069 |
| GO:0008016 | regulation of heart contraction | 1.465e-3 | 5/467 | 34/18671 | 5 | ENSG00000181800 ENSG00000203403 ENSG00000228113 ENSG00000259605 ENSG00000267280 ENSG00000267395 |
| GO:0010634 | positive regulation of epithelial cell migration | 1.905e-3 | 5/467 | 36/18671 | 5 | ENSG00000224086 ENSG00000232480 ENSG00000234899 ENSG00000270069 |
| GO:0001525 | angiogenesis | 5.795e-3 | 13/467 | 232/18671 | 13 | ENSG00000226252 ENSG00000227694 ENSG00000229153 ENSG00000232480 ENSG00000232973 ENSG00000236841 ENSG00000237512 ENSG00000239437 ENSG00000254481 ENSG00000255201 ENSG00000258168 ENSG00000268854 ENSG00000269124 ENSG00000267280 |
| GO:0007050 | cell cycle arrest | 5.994e-3 | 9/467 | 132/18671 | 9 | ENSG00000232480 ENSG00000239911 ENSG00000242808 ENSG00000244040 ENSG00000248636 ENSG00000249738 ENSG00000261924 ENSG00000267653 ENSG00000267194 ENSG00000271009 |
| GO:0007162 | negative regulation of cell adhesion | 7.394e-3 | 5/467 | 49/18671 | 5 | ENSG00000227935 ENSG00000231799 ENSG00000239437 ENSG00000255571 ENSG00000260788 |
| GO:0010923 | negative regulation of phosphatase activity | 8.050e-3 | 5/467 | 50/18671 | 5 | ENSG00000186594 ENSG00000225361 ENSG00000243902 ENSG00000256085 ENSG00000265222 ENSG00000265337 |
| GO:0035278 | miRNA mediated inhibition of translation | 8.268e-3 | 7/467 | 92/18671 | 7 | ENSG00000229989 ENSG00000247095 ENSG00000255248 ENSG00000255571 ENSG00000270069 |

##### 2.1.3 Pathway enrichment using predefined pathway gene sets

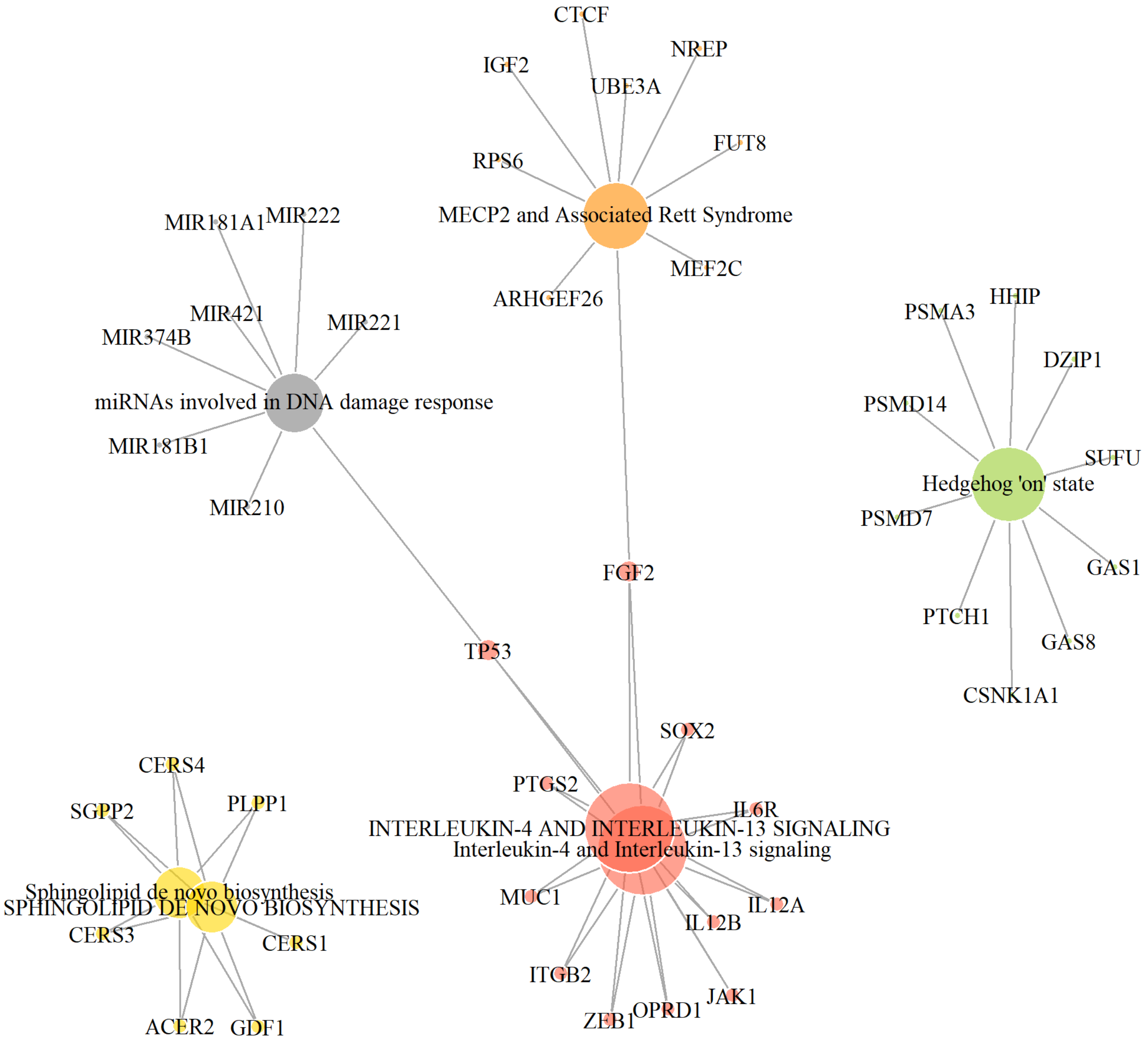

Figure S3: Interaction between the top 7 enriched pathways and the genes are depicted as a network. The size of the nodes represents the degree, number of edges that are incident to the vertex, and each color represents different clusters in the network.

Table S6: Neural disease related pathway enrichment results for the brain disease related ncRNA genes using customized pathway databases. If same pathway is detected in multiple database, we list the enriched pathway that have smaller p-adjusted value. The GeneRatio is computed by dividing the overlapping with the coding genes with the functional gene set to the number of all protein-coding genes within the input set neighbourhood. The BGRatio column represents the ratio of the number of genes found in the enriched GO term set to the size of the background gene set. The EGNo refers to the size of the overlap between the corresponding GO term gene set and the neighboring coding gene set. ncGeneList column contains ncRNA genes that are enriched with the corresponding GO-term.

| Pathway Term | Database | Pathway ID | Pvalue | GeneRatio | BGRatio | EGNo | ncGeneList |
|---|---|---|---|---|---|---|---|
| miRNAs involved in DNA damage response | WikiPathways | WP1545 | 3.489e-4 | 8/484 | 50/13604 | 8 | ENSG00000229989 ENSG00000230590 ENSG00000247095 ENSG00000262251 ENSG00000270069 ENSG00000271430 ENSG00000271533 |
| INTERLEUKIN-4 AND INTERLEUKIN-13 SIGNALING | REACTOME | R-HSA-6785807 | 5.793e-4 | 12/484 | 111/13604 | 12 | ENSG00000227039 ENSG00000228013 ENSG00000237036 ENSG00000240775 ENSG00000242808 ENSG00000244040 ENSG00000249738 ENSG00000262251 ENSG00000226852 ENSG00000226891 ENSG00000231064 ENSG00000273129 |
| SPHINGOLIPID DE NOVO BIOSYNTHESIS | REACTOME | R-HSA-1660661 | 7.325e-4 | 7/484 | 43/13604 | 7 | ENSG00000269694 ENSG00000271717 ENSG00000260912 ENSG00000261428 ENSG00000270127 |
| MECP2 and Associated Rett Syndrome | WikiPathways | WP3584 | 8.890e-4 | 9/484 | 71/13604 | 9 | ENSG00000180662 ENSG00000240775 ENSG00000240801 ENSG00000243069 ENSG00000246859 ENSG00000248309 ENSG00000224078 ENSG00000261386 ENSG00000273226 |
| Alzheimers Disease | WikiPathways | WP2059 | 9.268e-4 | 14/484 | 150/13604 | 14 | ENSG00000229989 ENSG00000245526 ENSG00000251034 ENSG00000253230 ENSG00000255248 ENSG00000255571 ENSG00000262251 ENSG00000235505 ENSG00000250377 ENSG00000257069 ENSG00000258399 ENSG00000261786 ENSG00000268401 |
| HEDGEHOG 'ON' STATE | REACTOME | R-HSA-56326846 | 0.0009 | 10/484 | 86/13604 | 10 | ENSG00000222019 ENSG00000232320 ENSG00000248890 ENSG00000257621 ENSG00000230551 ENSG00000271314 ENSG00000226237 ENSG00000239763 ENSG00000247400 ENSG00000272933 |
| SIGNALING BY HEDGEHOG | REACTOME | R-HSA-5358351 | 0.0012 | 13/484 | 137/13604 | 13 | ENSG00000222019 ENSG00000232320 ENSG00000248890 ENSG00000257621 ENSG00000230551 ENSG00000267009 ENSG00000271314 ENSG00000226237 ENSG00000236107 ENSG00000239763 ENSG00000247400 ENSG00000214765 ENSG00000272933 |

Table S7: Continued Table-Neural disease related pathway enrichment results for the brain disease related ncRNA genes using customized pathway databases. The GeneRatio is computed by dividing the overlapping with the coding genes with the functional gene set to the number of all protein-coding genes within the input set neighbourhood. The BGRatio column represents the ratio of the number of genes found in the enriched GO term set to the size of the background gene set. The EGNo refers to the size of the overlap between the corresponding GO term gene set and the neighboring coding gene set. ncGeneList column contains ncRNA genes that are enriched with the corresponding GO-term.

| Pathway Term | Database | Pathway ID | Pvalue | GeneRatio | BGRatio | EGNo | ncGeneList |
|---|---|---|---|---|---|---|---|
| Sphingolipid Metabolism (general overview) | WikiPathways | WP4725 | 0.0013 | 5/484 | 24/13604 | 5 | ENSG00000269694 ENSG00000271717 ENSG00000261428 ENSG00000270127 |
| Sphingolipid Metabolism (integrated pathway) | WikiPathways | WP4726 | 0.0016 | 5/484 | 25/13604 | 5 | ENSG00000269694 ENSG00000271717 ENSG00000261428 ENSG00000270127 |
| Prion disease pathway | WikiPathways | WP3995 | 0.0018 | 6/484 | 37/13604 | 6 | ENSG00000248309 ENSG00000255201 ENSG00000261386 ENSG00000235505 ENSG00000245812 ENSG00000255129 |
| Amyotrophic lateral sclerosis (ALS) | WikiPathways | WP2447 | 0.0021 | 6/484 | 38/13604 | 6 | ENSG00000251034 ENSG00000262251 ENSG00000267653 ENSG00000272163 ENSG00000235505 ENSG00000257069 ENSG00000267194 |
| Prader-Willi and Angelman Syndrome | WikiPathways | WP3998 | 0.0023 | 8/484 | 66/13604 | 8 | ENSG00000256164 ENSG00000259344 ENSG00000224078 ENSG00000262251 |
| AXON GUIDANCE | REACTOME | R-HSA-422475 | 0.0023 | 32/484 | 528/13604 | 32 | ENSG00000182000 ENSG00000196656 ENSG00000211451 ENSG00000227935 ENSG00000229153 ENSG00000229668 ENSG00000229848 ENSG00000231799 ENSG00000232320 ENSG00000236107 ENSG00000236432 ENSG00000237512 ENSG00000239437 ENSG00000242791 ENSG00000250107 ENSG00000251460 ENSG00000253389 ENSG00000255513 ENSG00000257621 ENSG00000260391 ENSG00000260778 ENSG00000272367 ENSG00000272944 ENSG00000273226 ENSG00000203589 ENSG00000217801 ENSG00000237943 ENSG00000239763 ENSG00000245750 ENSG00000255129 ENSG00000270172 ENSG00000273443 |

Table S8: Continued Table-Neural disease related pathway enrichment results for the brain disease related ncRNA genes using customized pathway databases. The GeneRatio is computed by dividing the overlapping with the coding genes with the functional gene set to the number of all protein-coding genes within the input set neighbourhood. The BGRatio column represents the ratio of the number of genes found in the enriched GO term set to the size of the background gene set. The EGNo refers to the size of the overlap between the corresponding GO term gene set and the neighboring coding gene set. ncGeneList column contains ncRNA genes that are enriched with the corresponding GO-term.

| Pathway Term | Database | Pathway ID | Pvalue | GeneRatio | BGRatio | EGNo | ncGeneList |
|---|---|---|---|---|---|---|---|
| Phosphodiesterases in neuronal function | WikiPathways | WP4222 | 0.0029 | 7/484 | 54/13604 | 7 | ENSG00000152931 ENSG00000213896 ENSG00000214244 ENSG00000225731 ENSG00000236460 ENSG00000242686 ENSG00000245958 ENSG00000270173 ENSG00000214765 |
| ACTIVATION OF NMDA RECEPTORS AND POSTSYNAPTIC EVENTS | REACTOME | R-HSA-442755 | 0.0033 | 8/484 | 70/13604 | 8 | ENSG00000217624 ENSG00000224086 ENSG00000239911 ENSG00000247572 ENSG00000248636 ENSG00000267009 ENSG00000269937 ENSG00000214765 |
| Synaptic signaling pathways associated with autism spectrum disorder | WikiPathways | WP4539 | 0.0091 | 6/484 | 51/13604 | 66 | ENSG00000239911 ENSG00000248636 ENSG00000224078 ENSG00000261924 ENSG00000260949 ENSG00000261044 |
| Signaling Pathways in Glioblastoma | WikiPathways | WP2261 | 0.0092 | 8/484 | 83/13604 | 8 | ENSG00000235369 ENSG00000255201 ENSG00000256164 ENSG00000258926 ENSG00000262251 ENSG00000267653 ENSG00000269937 ENSG00000237943 ENSG00000267194 |

##### 2.1.4 Comparison Between ASD Associated GO-terms and NoRCE Enrichment

Table S9: List of ASD-associated GO-terms that are identified by the NoRCE is provided for each analysis. ORatio column represents the observed ratio which is a fraction of the number of observed GO-terms to total number of identified GO-terms.

| Analysis | ORatio | GO-terms |
|---|---|---|
| Close-by Genes | 32/48 | GO:0070102 GO:0043542 GO:0007162 GO:0043542 GO:0007162 GO:0060045 GO:0034765 GO:0001525 GO:0008016 GO:0043536 GO:0045766 GO:0007050 GO:0035335 GO:0030148 GO:0006890 GO:0000209 GO:0051216 GO:0035278 GO:0032147 GO:0045669 GO:0016339 GO:0071407 GO:0008344 GO:0010923 GO:0045668 GO:0043087 GO:0008360 GO:0006470 GO:0006417 GO:0019233 GO:0006469 GO:0006887 |
| TAD based | 22/29 | GO:0060045 GO:0043536 GO:0008016 GO:0010634 GO:0001525 GO:0007050 GO:0007162 GO:0010923 GO:0035278 GO:0045766 GO:0035023 GO:0045669 GO:0035019 GO:0014068 GO:0034765 GO:0043687 GO:0006511 GO:0043087 GO:0032147 GO:0008360 GO:0006469 GO:0006816 |

#### 2.2 Case Study 3: Functional enrichment analysis with co-expression analysis

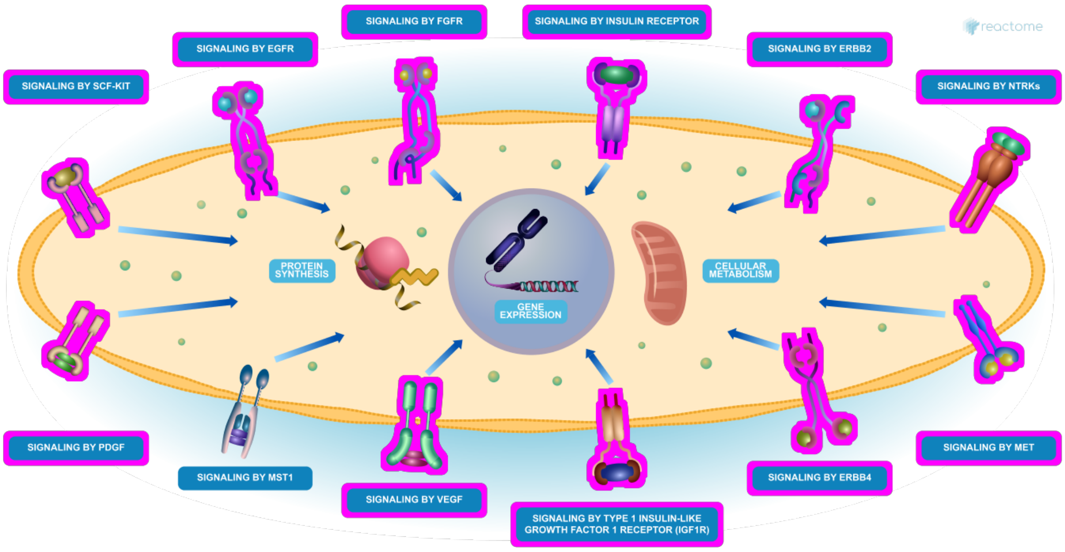

Figure S4: Reactome Diagram for the identified enriched pathway, Signaling by Receptor Tyrosine Kinases. For the given miRNA, coding genes that pass all filters for this analysis are flagged with purple color. Also, if those genes are annotated with another pathway in the diagram, positive pathways are also marked with purple color.
